# A Multicellular Network Mechanism for Temperature-Robust Food Sensing

**DOI:** 10.1101/815373

**Authors:** Dhaval S. Patel, Giovanni Diana, Eugeni V. Entchev, Mei Zhan, Hang Lu, QueeLim Ch’ng

**Author notes:** These authors contributed equally to this work (Dhaval S. Patel, Giovanni Diana, Eugeni V. Entchev).

## Abstract

Responsiveness to external cues is a hallmark of biological systems. In complex environments, organisms must remain responsive to specific inputs even as other internal or external factors fluctuate. Here we show how *Caenorhabditis elegans* can discriminate between food levels to modulate lifespan despite temperature perturbations. While robustness of fixed outputs has been described, our findings uncover a more complex robustness process that maintains food-responsiveness. This end-to-end robustness from environment to physiology is mediated by food-sensing neurons that communicate via TGF-β and serotonin signals to form a multicellular gene network. Mechanistically, specific regulations in this network change with temperature to maintain similar food-responsiveness in the lifespan output. Together, our findings provide a basis for gene-environment interactions and unveil computations that integrate environmental cues to govern physiology.

## Main Text

Robustness is the ability of a system to maintain its performance under perturbation (*1–3*). It is fundamental to biological systems, enabling organisms to thrive despite fluctuations in internal processes or external environments. Mechanisms for robustness exist in many precise and stereotyped processes, such as development (*1, 3*), circadian rhythms (*4*), and rhythmic neural activity (*5*), to produce invariant outputs despite internal variability in signalling activities or external perturbations in environmental conditions.

Unlike these stereotyped processes, much less is known about robustness in responsive processes in metazoans, where the ability to respond to one environmental cue is maintained despite fluctuations in a second factor that impact the same process. This form of robustness is crucial in complex natural environments where multiple factors can fluctuate independently. A rigorous understanding of robustness necessitates explicit definition of the biological parameter that is robust, and the perturbation that it is robust to (*1*). Here, we investigate how discrimination between food levels is robust to temperature in the nematode *C. elegans*.

Food and temperature affect lifespan in many species, including *C. elegans* (*6–17*). These effects can be observed when *C. elegans* are shifted to specific food and temperature levels during their reproductive period on day 2 of adulthood (Fig. 1A-B; Methods) (*18, 19*). At 20 °C, decreasing bacterial food concentration from *ad libitum* to starvation leads to local maxima and minima in lifespan, which plateaus to a maximum at the lowest food levels (*18*) (Fig. 1C). At a baseline food level (2×10^9^ bacterial cells/ml), decreasing temperature from 25 °C to 15 °C extends lifespan (Fig. 1D). Because *C. elegans* adopts a boom-and-bust lifestyle in temperate climates (*20*), these food and temperature ranges are consistent with fluctuations seen in its natural environment.

To determine how these environmental factors interact, we measured lifespan under 24 combinations of food and temperature (Fig. 1B). We then stratified the results by food or temperature to reveal the effect of temperature on food responsiveness and vice versa. At temperatures from 15 °C to 25 °C, the basic shape of the food-lifespan relationship remained similar even as increasing temperature reduced lifespan (Fig. 1E). Increased temperature also compressed the dynamic range of this food response, as measured by the difference between the highest and lowest mean lifespans across food levels (Fig. 1E). At food levels from starvation to *ad libitum*, increasing temperature generally reduced lifespan, but the dynamic range of the temperature response varied with food, and were highest at the lowest food levels (Fig. 1F).

**Figure 1.**
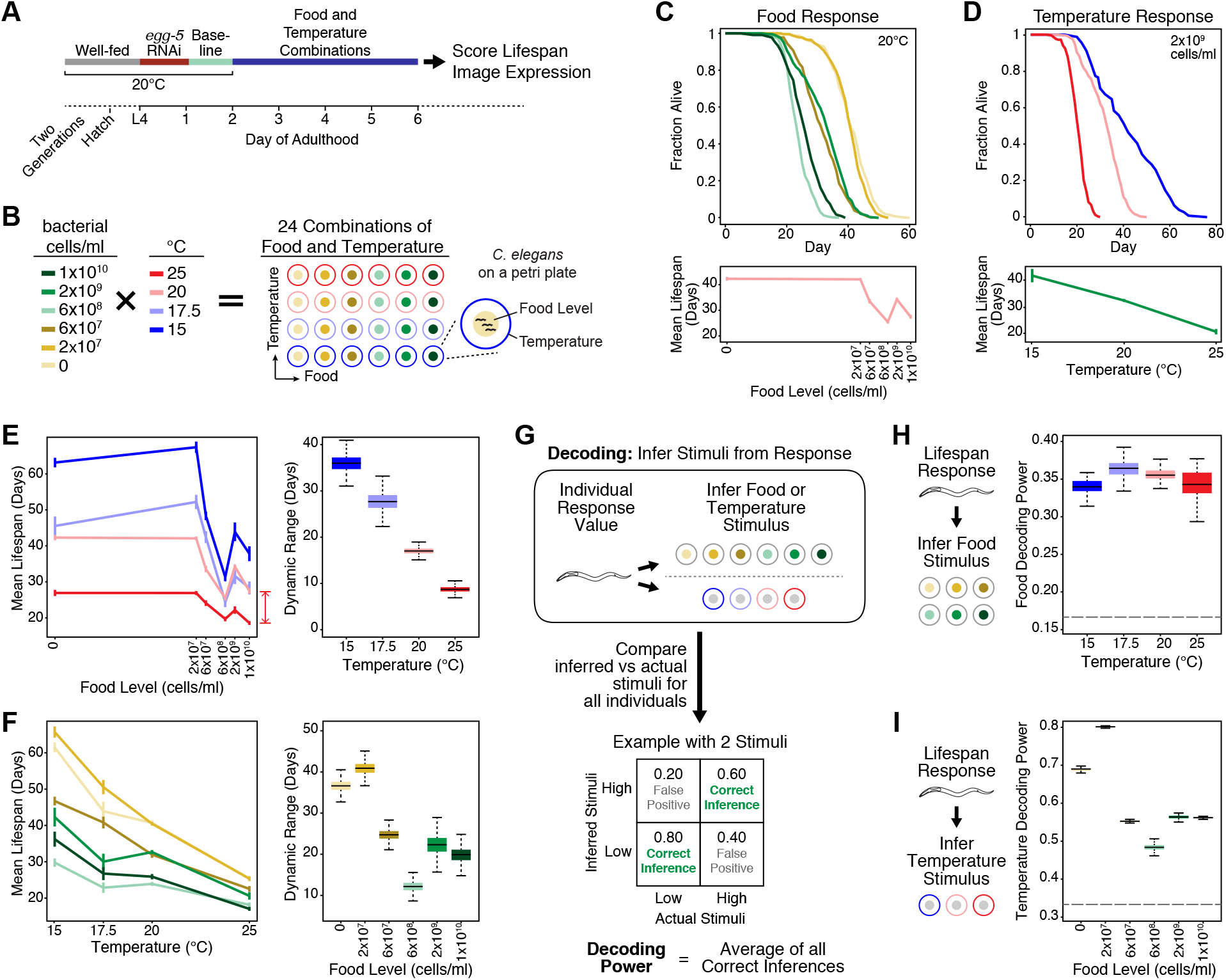
Combinatorial effects of food and temperature on *C. elegans* lifespan. **(A)** Food and temperature manipulations. Animals were shifted to specific food and temperature combinations after which their lifespan or gene expression levels were measured. **(B)** Legend for food and temperature combinations tested. **(C)** Effect of food levels on wild-type lifespan at 20 °C. Top: Kaplan-Meier Survival curves for each food level. Bottom: Mean lifespan across food levels. **(D)** Effect of temperature on wild-type lifespan at 2×10^9^ bacterial cells/ml. Top: Kaplan-Meier Survival curves for each temperature. Bottom: Mean lifespan across temperature. **(E)** Effect of temperature on food responsiveness of lifespan. Left: Mean lifespan across food levels at different temperatures. Right: Dynamic range of food responsiveness as indicated by red bar at each temperature. **(F)** Effect of food on temperature responsiveness of lifespan. Left: Mean lifespan across temperature at different food levels. Right: Dynamic range of temperature responsiveness. **(G)** Scheme for decoding analysis to quantify discrimination. Examples of inferred frequencies are provided to illustrate the analysis output. See text and supplementary methods for details. **(H)** Food decoding power at different temperatures based on lifespan responses. **(I)** Temperature decoding power at different food levels based on lifespan responses. Bayesian estimates are shown for all mean lifespans and dynamic ranges (see Methods and Table S1). Error bars in all line plots in c-f indicate standard deviations. Bayesian distributions in (E, F, H, and I) are depicted by boxplots. Dotted lines in (H-I) indicate decoding power from random chance alone.

This ability to produce different lifespans under different environmental conditions implies that *C. elegans* can discriminate among them. We used decoding analysis (*21*) to quantify how well environmental conditions can be discriminated based on the corresponding distributions of lifespan or gene expression. This method has been successfully applied to gene expression, biochemical activities, and physiological responses (*18, 22*), and allows us to accommodate noisy responses and non-linear stimuli-response relationships. Using this approach (Fig. 1G; Methods), we first infer the most likely stimulus for a given response, using prior knowledge of the response distributions under different stimuli. Next, we compared the inferred versus actual stimuli to determine the frequency of correct and incorrect inferences under each stimulus, generating a matrix of inference patterns depicting how well different stimuli are distinguishable from each other (Fig. 1G and S1). Finally, we use the average frequency of correct inferences, which we term decoding power, to summarize the discriminatory performance. Higher decoding power is associated with a stronger coupling between stimuli and response, indicating better discrimination and hence better performance.

For each temperature, we used decoding analysis to quantify discrimination between food levels that produces different lifespans. Remarkably, food decoding power in wild-type animals remained constant from 15 °C to 25 °C (Fig. 1H), despite the large changes in the absolute lifespan values (Fig. 1E). Some food levels were discriminated well and others poorly, and this inference pattern underlying the average decoding power was also preserved across this 10 °C range (Fig. S1A). Thus, discrimination between food levels is robust to temperature. This robustness is specific: although discrimination between temperatures is relatively strong, the performance fluctuates considerably as food levels are changed (Fig. 1I and S1B). This robustness represents a novel interaction between food and temperature during lifespan modulation; it is biologically significant because it spans from environmental input to physiological output.

To understand how robustness arises, we assessed the role of food sensing pathways. TGF-β and serotonin are signals in conserved nutrient sensing pathways that regulate diverse physiological processes in many species (*23–30*). In *C. elegans*, disrupting these pathways by gene deletions of *daf-7* [encodes TGF-β (*23*)] or *tph-1* [encodes tryptophan hydroxylase, the rate-limiting enzyme in serotonin synthesis (*25*)] ablates most of the ability to modulate lifespan in response to food (*18, 31*). Since these genes represent a major link between food and lifespan, we assessed their contributions to robustness in food level discrimination at different temperatures (Fig. 2). At all temperatures, loss of *tph-1* and *daf-7* attenuated both increases and decreases in lifespan due to food, with the most severe attenuation observed in the double mutant (Fig. 2). This attenuation was most obvious under starvation and at 6×10^8^ bacterial cells/ml, where the respective food-dependent lifespan extension and reduction occurred with smaller magnitudes in the single and double mutants compared to wild-type (Fig. 2A).

**Figure 2.**
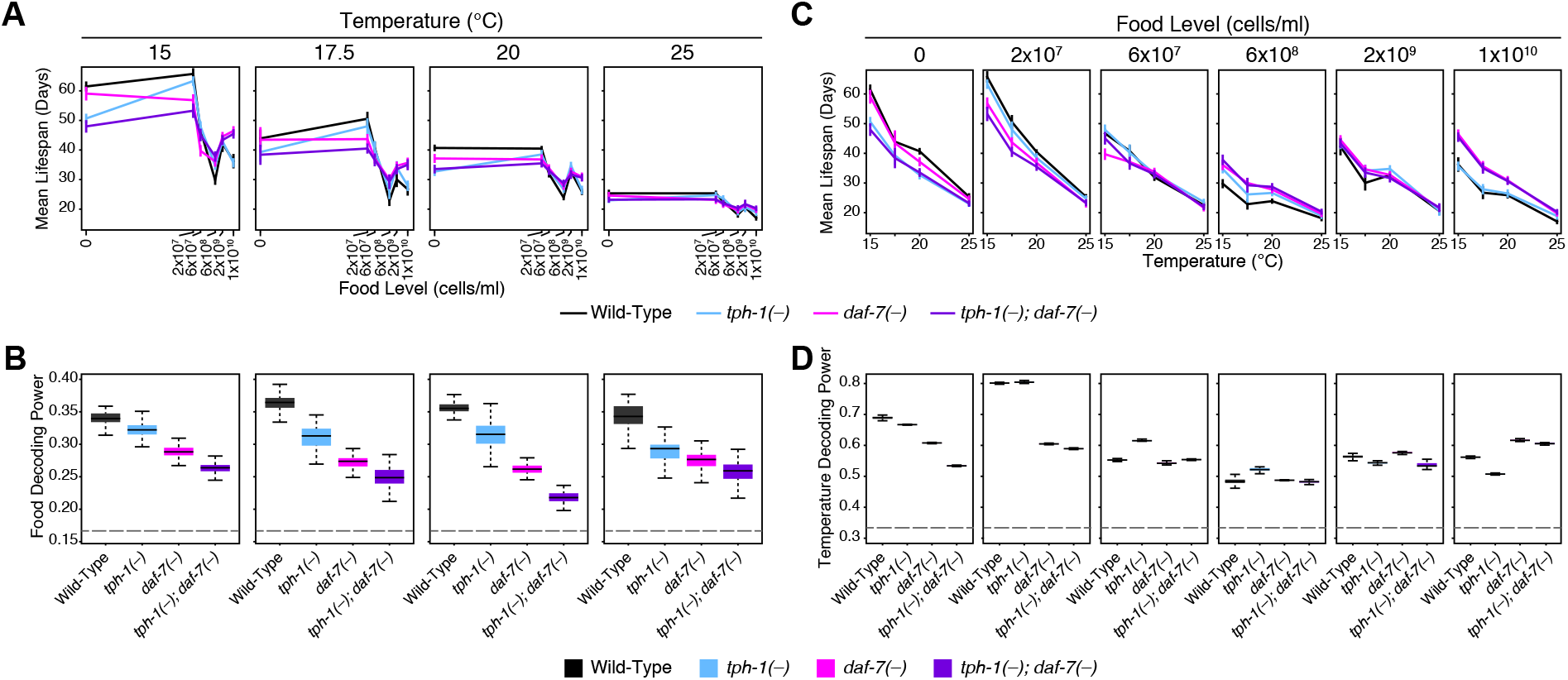
*tph-1* and *daf-7* are required for temperature-robust food responsiveness in lifespan. **(A)** Mean lifespan for each genotype as a function of food levels under different temperatures. Loss of *tph-1* and *daf-7* attenuate food responsiveness bidirectionally at all temperatures. **(B)** Mean lifespan for each genotype as a function of temperature at each food level. **(C)** Food discrimination based on the lifespan response is reduced in *tph-1*(*-*) and *daf-7*(*-*) at all temperatures. **(D)** Effect of *tph-1*(*-*) and *daf-7*(*-*) on temperature discrimination based on lifespan. Bayesian estimates are shown for all mean lifespans (see Methods and Table S1). Error bars in all line plots (A, C) indicate standard deviations. Bayesian distributions (B, D) are depicted by boxplots. Dotted lines in (B, D) indicate decoding from random chance alone.

Decoding analysis of the lifespan responses revealed that at all temperatures tested, food decoding power was reduced in both *tph-1*(*-*) and *daf-7*(*-*) mutants, with more severe effects in the double mutant (Fig. 2B). While the pattern of correct and incorrect inferences in wild-type were largely stable to temperature, these patterns became more temperature-sensitive in *tph-1*(*-*) and *daf-7*(*-*) single and double mutants (Fig. S1A), implying that unlike wild-type, the mutants were attuned to different food levels at different temperatures. Thus, this effect on food decoding power is not a trivial consequence of *tph-1*(*-*) and *daf-7*(*-*) mutants being unable to sense food. Instead, they indicate that *tph-1* and *daf-7* contribute to the robustness of food discrimination to temperature perturbations.

In contrast, *tph-1* and *daf-7* did not mediate robustness of temperature discrimination to food levels. Loss of *tph-1* and/or *daf-7* enhanced or reduced the effects of temperature depending on the food level (Fig. 2C), leading to corresponding increases or decreases in temperature decoding power as a function of food (Fig. 2D). Taken together, we conclude that the effects of *tph-1* and *daf-7* on robustness are specific to one functional parameter (food level discrimination) under one perturbation (temperature).

Robustness could be achieved in two general ways: either by resisting the perturbation and remaining unchanged, or by adapting and compensating. To distinguish between these possibilities, we examined *tph-1* expression in NSM and ADF sensory neurons and *daf-7* expression in ASI sensory neurons (Fig. 3A), where these genes are primarily expressed (*23, 25, 32*), and act from, to modulate lifespan (*18, 33*). In these food-sensing neurons (*18, 34–36*), the expression levels of *tph-1* and *daf-7* are regulated by food levels to affect lifespan, thereby providing a representation of food abundance with a functional output (*18, 33*). Cross- and self-regulation among these genes in their respective cells indicate that they act in a gene network distributed over multiple cells (*18, 25, 28*). We therefore characterized wild-type gene expression in this network under a systematic combination of food levels and temperatures (Fig. 3B), using quantitative high-throughput imaging to measure expression of validated single-copy transcriptional reporters for *tph-1* and *daf-7* at the single-cell level (*18, 37, 38*).

**Figure 3.**
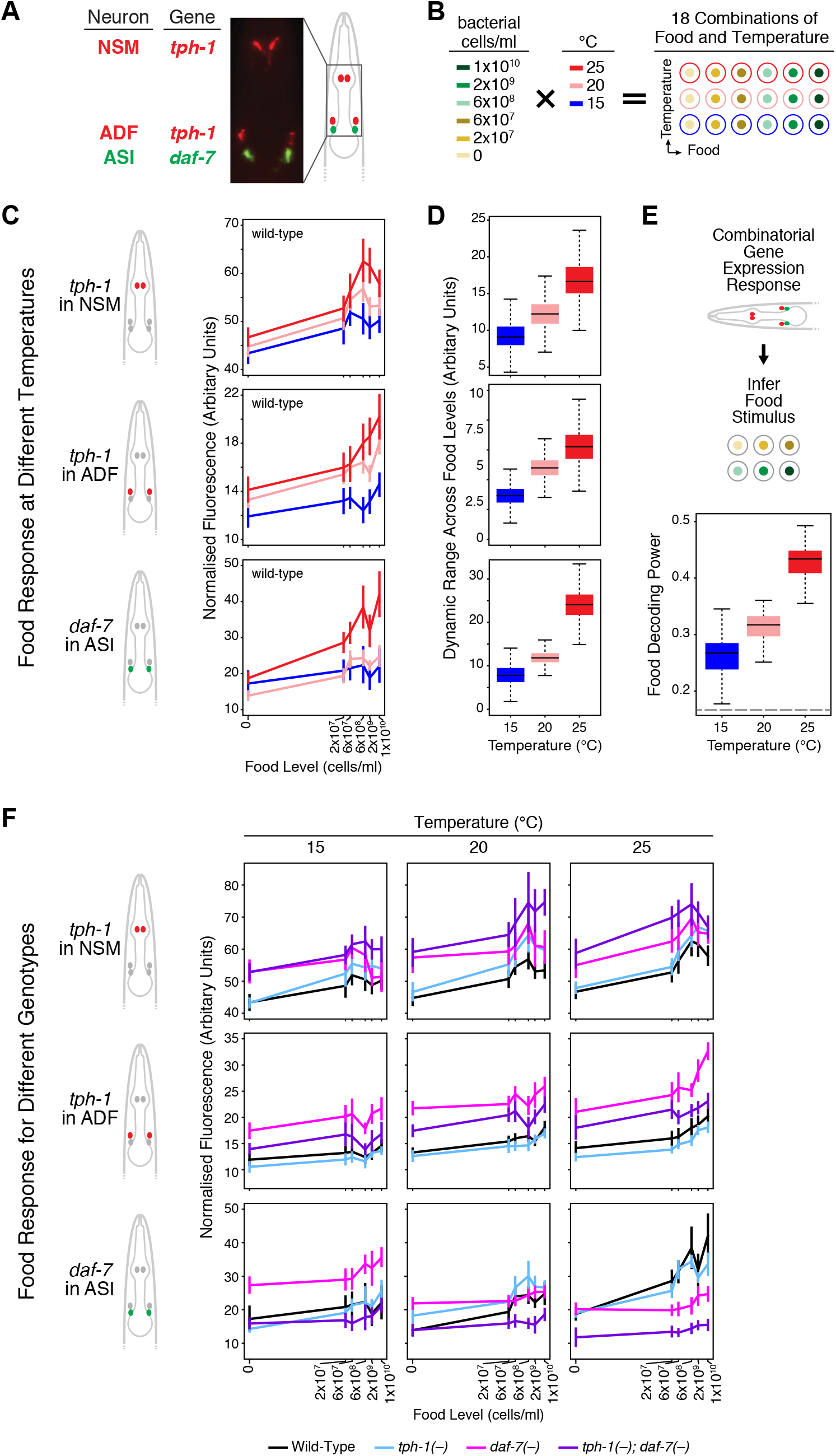
High-throughput gene expression analysis of *tph-1* and *daf-7*. **(A)** Transcriptional reporters showing *tph-1* expression in NSM and ADF neurons, and *daf-7* expression in ASI neurons. **(B)** Legend for 18 food and temperature combinations tested. **(C)** Wild-type expression of *tph-1* in NSM and ADF, and *daf-7* in ASI as a function of food. Each line corresponds to a different temperature. Diagrams on the left denote gene and cell. **(D)** Dynamic range of food responsiveness in expression of *tph-1* in NSM and ADF, and *daf-7* in ASI at different temperatures. **(E)** Wild-type food decoding power at different temperatures based on the combinatorial expression of *tph-1* and *daf-7* in these three cells. The data in (C-E) is plotted as a function of temperature in Figure S2. **(F)** Expression of *tph-1* in NSM and ADF, and *daf-7* in ASI as a function of food at different temperatures in single and double mutants of *tph-1* and *daf-7*. To visualize this expression data as a function of temperature at different food levels, see Figure S3. Bayesian estimates are shown for all mean expression values and dynamic ranges (see Methods and Table S2). Error bars denote 90% confidence intervals. Dotted line in (E) indicates decoding from random chance alone.

In wild-type animals, expression of *tph-1* in NSM and ADF or *daf-7* in ASI, based on these reporters, show distinct response profiles as a function of food, reflecting their food encoding properties (*18*) (Fig. 3C). While these food response curves largely retained their shape from 15-25 °C, expression levels generally increased with higher temperatures (Fig. 3C and S2). Additionally, the dynamic range of gene expression for both genes in all three sets of cells also increased with temperature, suggesting that these gene activities are more responsive to food levels (Fig. 3D).

To quantify discrimination at the level of gene expression, we used decoding analysis. Having simultaneously imaged all three cells in each animal, we could infer stimuli based on the combinatorial expression values of *tph-1* and *daf-7* in NSM, ADF, and ASI (Fig. 3E). This analysis revealed that the food decoding power of this combinatorial gene expression increased with temperature (Fig. 3E). In other words, these cells could distinguish between food levels better as temperature was increased, consistent with the increased dynamic range of their food responses (Fig. 3D).

These results suggest that robustness to temperature is achieved by changes in the underlying multicellular gene network, whereby the network adapts to compensate for temperature rather than remain unchanged. As temperature increased from 15 to 25 °C, the magnitudes of food-dependent lifespan changes were compressed and the effects of *tph-1* and *daf-7* on lifespan were reduced (Fig. 1E and 2A). Increasing the dynamic range of food-responsive *tph-1* and *daf-7* expression (Fig. 3D) heightens the food-dependent difference in gene activity and therefore compensates at least in part for the reduced impact of these genes. This increased decoding power from the combinatorial gene expression revealed that the discriminatory ability of the multicellular gene network is plastic and temperature dependent. This finding also raised the question of how the performance of such a higher-order function as discrimination, which involves mapping multiple stimuli to corresponding responses, could be modulated.

We previously identified modulatory regulation among *tph-1* and *daf-7* at 20 °C using transcriptional reporters to measure *tph-1* and *daf-7* expression levels in NSM, ADF, and ASI in single and double mutants of these genes at different food levels (*18, 31*). To understand how these interactions are affected by temperature, we analyzed the effects of *tph-1* and *daf-7* mutations on the expression of these reporters in their respective cells under more extensive food and temperature combinations (Fig. 3B and F). These results revealed how regulations among *tph-1* and *daf-7* change as a function of food and temperature. First, mutations in *tph-1* and *daf-7* could affect expression of *tph-1* and *daf-7* in all three cells, indicating that *tph-1* and *daf-7* act in a highly inter-connected network with extensive cross- and self-regulation. Second, *tph-1* and *daf-7* act separately but interact to set gene expression levels. In all three cells, the expression phenotypes of *tph-1*(*-*)*; daf-7*(*-*) double mutants largely differed from each of the single mutants (Fig. 3F and S3). Furthermore, comparisons between *daf-7*(*-*) single mutants and *tph-1*(*-*)*; daf-7*(*-*) double mutants show that *tph-1*(*-*) mutations have a more prominent effect in the *daf-7*(*-*) background (Fig. 3F and S3). Third, gene-environment interactions were extensive, as food and temperature modified the effects of *tph-1*(*-*) and *daf-7*(*-*) mutations. Using *daf-7* expression in ASI as an example (Fig. 3F and S3 bottom row), loss of both *tph-1* and *daf-7* had a greater effect on *daf-7* expression in ASI at higher food levels. Also, loss of *daf-7* increased *daf-7* expression in ASI at 15 °C but decreased it at 25 °C, indicating a temperature dependence in *daf-7* self-regulation. Such temperature-dependent phenotypes suggest that the network configuration changes with temperature. Because *tph-1* and *daf-7* mediate robustness to temperature (Fig. 2), our results suggest that these genes adopt different network configurations at different temperatures to produce similar food discrimination.

To understand the temperature-dependent changes in this *tph-1*/*daf-7* network, we used *in silico* modelling to disentangle its complex interactions (Fig. 4, Methods). We modelled this network as three interconnected nodes (*tph-1* in NSM, *tph-1* in ADF, and *daf-7* in ASI). Since loss of *tph-1* and *daf-7* independently affects expression of both genes in all cells, there are a total of nine effective connections that represent the net regulation of one node by another (Fig. 4A). Each connection could represent positive or negative regulation, resulting in 2^9^ (= 512) possible networks. Conventionally, if loss of gene A leads to increased expression of gene B, one would infer that the best fit model is gene A inhibiting gene B. We embedded this type of logic within the computational model to describe the net regulatory effects that were consistent with the observed gene expression phenotypes.

**Figure 4.**
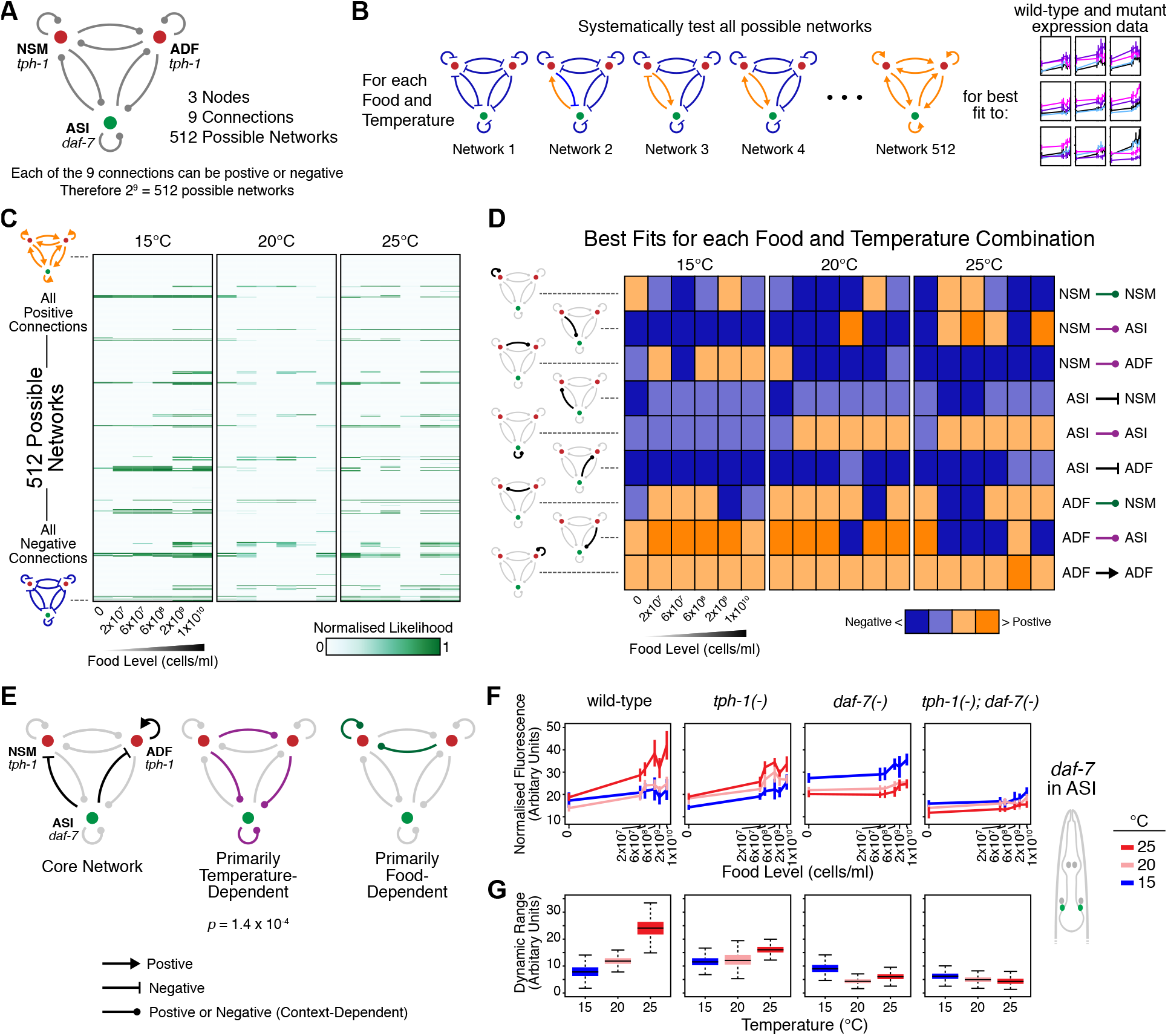
Temperature-dependent changes in the multicellular *tph-1*/*daf-7* network. **(A)** Models of a multicellular network comprising *tph-1* in NSM and ADF, and *daf-7* in ASI. **(B)** Identifying best-fit models of the network at each food and temperature combination, based on expression data from Figure 3. **(C)** Heat map summarizing goodness of fit for all 512 models across 18 food-temperature combinations. Each row corresponds to a specific network model, each column is a food-temperature combination. **(D)** Summary of best fit models for each food-temperature combination. Each edge is categorized as positive or negative regulation and as strong or weak. Each row denotes a specific connection (left and right labels); each column indicates a food-temperature combination. **(E)** Some regulatory connections are invariant (Core Network), whilst others change between positive and negative regulation depending on the environment. *p* indicates significance of the temperature-dependent configurations from bootstrapping. **(F)** Role of temperature-dependent connections on *daf-7* expression in ASI revealed by loss of *tph-1* or *daf-7*. Bayesian estimates for *daf-7* expression in ASI are shown; error bars indicate 90% confidence intervals. **(G)** Temperature-dependent increase in the food-responsive dynamic range of *daf-7* expression in ASI requires temperature-dependent connections that converge on *daf-7* expression in ASI. Bayesian distributions of dynamic range are indicated by boxplots.

We then characterized the effects of the environment in several steps. First, considering only one food-temperature combination, we determined which of the 512 possible network models provided the best fit for the expression of both genes in all three cells across all four genotypes (Fig. 4B). Next, we repeated these fits across each of the 18 food-temperature combinations (Fig. 4B) and visualized how well each network configuration explains the gene expression phenotypes at each environmental combination (Fig. 4C). Network configurations that fit well were sparse, indicating that our data provided sufficient constraints. Many networks that fit well were similar, as shown by the repeated spacing between the good fits within each column, which differed by the sign of only one connection. We then visualized the best-fit network for each food-temperature combination (Fig. 4D) by categorizing each connection as positive or negative and as strong or weak, which revealed stable and variable connections. For example, the computational analysis suggested that *daf-7* in ASI negatively regulates *tph-1* expression in NSM and ADF across all food and temperature conditions (Fig. 4D, 4^th^ and 6^th^ rows), consistent with experimental data showing that *daf-7*(*-*) mutants possess higher *tph-1* expression in NSM and ADF than wild-type in all conditions tested (Fig. 3F). In this manner, we used model-fitting as an analytical approach to identify putative regulatory interactions that were stable as well as those that primarily varied with temperature or food (Fig. 4E).

These computational analyses pointed to the regulatory interactions that changed sign with temperature, three of which converged on *daf-7* expression in ASI (Fig. 4E, center). This convergence was notable because the food-responsive dynamic range of *daf-7* expression in ASI showed the greatest temperature-dependent increase among all three cells (Fig. 3D), leading us to focus on the subset of our experimental data (Fig. 3F, S3, and S4) involving the regulation of *daf-7* expression in ASI (Fig. 4F). First, we considered the role of the cross-regulation from *tph-1* in NSM and ADF to *daf-7* in ASI by examining experimental data from the *tph-1*(*-*) mutant where these connections are ablated. In wild-type animals, the dynamic range of food-responsive *daf-7* expression in ASI increased with temperature (Fig. 3D, 4F-G). This increase in dynamic range was impaired in *tph-1*(*-*) mutants (Fig. 4G), indicating that *tph-1* is required for increased food-responsiveness of *daf-7* expression in ASI at higher temperatures.

Next, we considered the role of *daf-7* self-regulation by examining the experimental data from the *daf-7*(*-*) mutant. In wild-type, expression levels of *daf-7* in ASI increased with higher temperature (Fig. 4F). In *daf-7*(*-*) mutants, this temperature effect was reversed: *daf-7* expression in ASI decreased with higher temperature (Fig. 4F). Thus, the increased expression at higher temperature was not simply due to thermodynamics; instead, it is genetically controlled. This autoregulatory effect of *daf-7* was not due to indirect feedback through *tph-1* because this inverted response to temperature was independent of *tph-1*: it was not observed in *tph-1*(*-*) mutants and still occurred when *daf-7*(*-*) was mutated in the *tph-1*(*-*) background [compare *tph-1*(*-*) to *tph-1*(*-*)*; daf-7*(*-*) in Fig. 4F]. *daf-7* self-regulation also contributed to increasing dynamic range of *daf-7* expression in ASI with higher temperature, and loss of both *tph-1* and *daf-7* abolished the temperature-dependent increase in the dynamic range of food-responsive gene activity (Fig. 4G).

These complementary results from computational analysis and experiments independently revealed roles of specific connections in the *tph-1*/*daf-7* network that vary with temperature. These temperature-dependent connections transform the way food is encoded by *daf-7* expression in ASI, from a representation of food abundance with a constant dynamic range into one where the dynamic range increases with temperature (Fig. 4G). These transformations reflect specific computations performed by specific connections to compensate for the reduced impact of *tph-1* and *daf-7* on lifespan as temperature increases, thereby providing a mechanism that contributes to temperature-robust food discrimination at the level of lifespan outputs (Fig. 1H).

This end-to-end robustness reflects a biologically relevant link from environment to physiology, and explains the need for food- and temperature-responsiveness, as well as robustness to temperature. Lifespan extension at low food levels enables *C. elegans* to reproduce much later when food becomes plentiful (*10, 39*). It is therefore advantageous to maintain such phenotypic plasticity within the reproductive temperature range. This temperature-robust feature requires temperature responsiveness in the *tph-1*/*daf-7* network to elicit compensatory changes (Fig. 4E-G), demonstrating that plasticity to one factor is required for robustness to a second factor.

Much work on robustness has focused on processes that generate a stereotyped output such as the development of specific anatomical structures or the production of precise rhythms (*1–5*). Here, we provide new insights into temperature-robustness of a responsive process where different food levels must be distinguished to produce different lifespans. This type of robustness in a higher-order function only became apparent when we used decoding analysis to quantify discrimination.

Feedback, buffering, and redundancy are robustness mechanisms that enable biological systems to compensate against perturbations (*1, 2*). These mechanisms have been implicated in many temperature-robust processes, such as buffering by microRNAs during development (*1*), feedback regulation of ion channels in neural circuit activity (*40*), as well as intercellular coupling at the organismal level and allosteric feedback at the molecular level in circadian oscillations (*4, 41*). Robustness is also tied to degeneracy, the ability of different components to impart the same function (*42*). In degenerate systems with distinct components that are sensitive to different conditions, robustness is ensured because some components remain functional when others are perturbed (*5, 43, 44*). Our data suggests that *tph-1*/*daf-7* network shows a different implementation, where degeneracy is induced by a perturbation that alters the configuration of its components. In turn, this induced degeneracy produces similar discriminatory performance under different conditions to generate robustness to the perturbation.

Three principal implications arise from our finding that different environments induce different network configurations to produce the same performance. First, biological networks in general, and gene regulation in particular, are not static. It will be particularly intriguing to understand the context-dependent organization of many biological networks to learn how their changes relate to their functions in complex environments. Second, robustness can be genetically controlled. We reveal that serotonin and TGF-β signals impinge on the robustness of food-sensing to temperature. These results align with precedents uncovering important roles for neuromodulators in temperature-robust rhythmic neural activity and thermosensory behavior (*43, 45*). Third, gene-environment interactions are a by-product of induced degeneracy, where under different conditions, network components possess different functions and therefore exhibit different loss-of-function phenotypes. If induced degeneracy is a common robustness mechanism, we might expect studies of gene-environment interactions to highlight robustness in additional areas of biology.

## Materials and Methods

### *C. elegans* Strains and General Culture

All strains of *C. elegans* were cultured using standard conditions (*46*). The following strains were used for the lifespan analyses: N2 (wildtype), QL101 *tph-1(n4622) II*, QL282 *daf-7(ok3125) III*, and QL300 *tph-1(n4622) II; daf-7(ok3125) III*. For quantitative imaging analyses the following strains were used: QL196 *drcSi61*[*tph-1p::mCherry, Cbr-unc-119(+)*] *I; drcSi7*[*daf-7p::Venus, Cbr-unc-119(+)*] *II*, QL404 *drcSi61*[*tph-1p::mCherry, Cbr-unc-119(+)*] *I; tph-1(n4622) drcSi7*[*daf-7p::Venus, Cbr-unc-119(+)*] *II*, QL402 *drcSi61*[*tph-1p::mCherry, Cbr-unc-119(+)*] *I; drcSi7*[*daf-7p::Venus, Cbr-unc-119(+)*] *II; daf-7(ok3125) III*, and QL435 *drcSi61*[*tph-1p::mCherry, Cbr-unc-119(+)*] *I; tph-1(n4622) drcSi7*[*daf-7p::Venus, Cbr-unc-119(+)*] *II; daf-7(ok3125) III*.

### Lifespan

Lifespans were carried out as previously described (*18, 19*) (Fig. 1A). Briefly, starting with starved animals, animals were raised for two generations on standard NGM culture plates (*46*) seeded with the live OP50 at the baseline temperature of 20 °C. The L4-stage progeny of the F2 generation were then transferred to RNAi plates seeded with bacteria expressing double-stranded RNA (dsRNA) targeting the *egg-5* gene, which is critical for egg-shell formation (*47, 48*), for 24 hours at 20 °C. This short period of exposure to *egg-5* RNAi is sufficient to prevent progeny production in the majority of animals throughout their reproductive period (*18, 19*). This step is especially necessary for *tph-1*(*-*) and *daf-7*(*-*) mutants, as well as the double mutant of these genes, as they have an impaired egg-laying phenotype which results in the premature death of the adult worm from matricide. After the 24-hour *egg-5* RNAi exposure the 1-day old adults were moved to NGM culture plates supplemented with the antibiotics streptomycin and carbenicillin (NSC) seeded with our baseline food level, antibiotic-inactivated OP50 at a concentration of 2×10^9^ cells/ml (*18, 19*), for 24 hours at 20 °C. All manipulations away from the baseline temperature and food level occurred on day 2 of adulthood (Fig. 1A). In total, we examined lifespan across 4 different temperatures and 6 different food levels (Fig. 1B). Animals at each temperature were manually transferred to new NSC plates seeded with the appropriate food concentration according to the schedule laid out in Patel et al (*19*). Animals were scored for death at every transfer point and then daily after the final transfer point for each temperature. Death was assessed as the failure to detect movement in response to a gentle prod with a wire pick. Part of the raw data for lifespans at 20°C was obtained from Entchev et al (*18*).

### Quantitative Fluorescence Imaging

Gene expression activity of *daf-7* and *tph-1* at single-cell and single-worm resolution were quantified based on the fluorescence intensity of *daf-7::Venus* and *tph-1::mCherry* transcriptional reporters as previously described (*18*). These reporters were integrated in single-copy and contain endogenous 5’ and 3’ sequences up to the next adjacent open reading frame; and their construction and validation were detailed in Entchev et al (*18*). Multiple labs have validated that transcriptional reporter fluorescence correlates with downstream activity in these respective pathways (*18, 23, 24, 33*).

For quantitative imaging, strains were cultured using a procedure similar to that in lifespan studies (*18, 19*). Animals were initially raised on large NGM plates (10 cm diameter) for two generations. Once the F2 animals reached reproductive adulthood, they were washed from the plates using S-basal supplemented with streptomycin and then subjected to a sodium hypochlorite treatment to break them apart and release their eggs (*46*). The eggs were then deposited on to new large NGM plates seeded with live OP50 and left to hatch and grow at 20 °C till the animals reached the L4-stage. The *daf-7(ok3125)* mutation causes animals to constitutively enter a transient dauer arrest (*18, 19*). For this reason, strains harboring *daf-7(ok3125)* were prepared 24 hours before strains without this mutation, to ensure sufficient numbers of animals exited dauer and reached the L4-stage. Once animals reached the L4-stage, they were collected by washing and distributed on to large RNAi plates seeded with *egg-5*-dsRNA expressing bacteria at a density of ∼100 worms per plate. Animals were exposed to *egg-5* RNAi for 1 day at 20 °C before being moved to large NSC plates seeded with our baseline food level for a further 24 hours also at 20 °C. As with the lifespan studies, all temperature and food manipulations were initiated on day 2 of adulthood. Worms at each temperature were transferred by washing to new large NSC plates seeded with the appropriate food concentration according to the schedule laid out in Patel et al (*19*). Animals were harvested for imaging upon reaching their sixth day of adulthood.

Imaging studies were performed on the microfluidic system previously described (*18*). Briefly, our imaging system consists of a custom two-layer microfluidic device (*37, 49*) constructed from polydimethylsiloxane (PDMS) using standard multilayer soft lithographic techniques (*50*) and bonded to a glass coverslip. Animals were loaded into the imaging area of the device using pressure driven flow controlled by on-chip valves that reside in the first layer of the device. The second layer of the device cooled the imaging area down to ∼4 °C, which immobilizes the worms during image capture. Images of the head of animal were captured as Z-stacks in 2 μm steps using a Nikon Ti-Eclipse inverted scope with a 40x (1.3 NA) oil objective on a Hamamatsu Orca R2 camera. The fluorescence intensities of the Venus and mCherry reporters was captured simultaneously using an Optosplit II system. Operation of worm-loading on the microfluidic system and subsequent image acquisition was automated through the use of a custom LabView code. Images were stored locally for offline processing using a custom MATLAB script that automatically assigned cell ID and extracted fluorescent intensities for the ASI, ADF, and NSM neurons in each stack. The code for quantification of the fluorescence signals from these neurons is publicly available on GitHub (https://github.com/meizhan/SVMelegans). Part of the raw data for gene expression at 20°C was obtained from Entchev et al (*18*).

### Statistical Analysis

To account for biological and experimental variability between and within experiments, we adopted a Bayesian framework to analyze the lifespan and gene expression data. We used hierarchical models to describe these sources of variability, and Bayesian inference to estimate features such as mean lifespan and expression levels. Throughout these analyses we used non-informative (flat) priors for all model parameters to minimize their effect on posterior distributions.

#### Lifespan

We modelled lifespan as a Weibull distribution with trial-dependent scale and shape parameters to accommodate batch variability. To estimate the average lifespan for each environmental condition with censoring using a data augmentation technique, we used the RJAGS package in R (*51*).

#### Gene Expression

To estimate *daf-7* and *tph-1* expression levels in their respective cells across all our experiments, we designed a Bayesian approach to incorporate all sources of experimental and biological variability within a unified hierarchical model (Fig. S6A). This model allows us to simultaneously quantify the effect of trial-to-trial variability, population variability, changes in LED illumination, and levels of correlation among *tph-1* and *daf-7* in ADF, ASI, and NSM. To correct for changes in LED illumination, we periodically imaged fluorescent bead standards in both red and green channels and used linear regression of these measured values over time to obtain normalization factors for each experiment. The posterior distributions of these regression coefficients were calculated as the product of an inverse gamma distribution associated with the time-independent variance in bead fluorescence and the multivariate normal distribution of their fluorescence (*52*). For each food and temperature condition, the unnormalized mean batch fluorescence 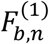 for cell *n* of batch *b* imaged at time *τ_b_* is expressed as:

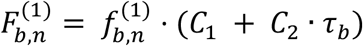

where *C_1_* and *C*_2_ are the linear regression coefficients for the relevant imaging channel modelled statistically according to the posterior distributions conditional to bead measurements, and 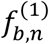 is the batch-level normalised fluorescence for each neuron. Normalized fluorescence levels for all cell is modelled with a multivariate normal distribution as described in Fig. S6A. We implemented this model using the RJAGS package in R (*51*) which performs Monte Carlo Gibbs sampling and provides the posterior distribution of all model variables given the raw data, allowing us to estimate the true fluorescence values and its uncertainty. The Gibbs sampler returns a list of values (Markov chains) for each model variable distributed according to their corresponding posterior distributions. The convergence of the sampler was estimated by requiring the convergence of 8 independent Markov chains to the same set of model parameters. Custom R scripts and data used in the above analyses are publicly available on GitHub (https://github.com/giovannidiana/TRFS).

### Decoding Analysis

To measure discrimination between different food and temperature conditions based on either expression or lifespan (Fig. S6B), we first expressed the probability of each response under each environmental condition based on the Bayesian analysis above. The maximum-likelihood decoder attributes each response to the environmental stimuli that maximizes the conditional probability. By applying the maximum-likelihood decoder to the data from all individuals, we can calculate the confusion matrix (Fig. S1 and S5) by counting how many times a response measured under a given environment is decoded as the correct environment, or as one of the incorrect environments. The closer the confusion matrix to a diagonal matrix, the more informative the response is about the state of the environment (Fig. S6B). To apply the maximum-likelihood decoder to our gene expression models, we estimated the probability distribution of the normalized expression levels for all environmental conditions. Because the normalization uncertainty is small compared to population and trial-to-trial variability, we approximated these distributions as multivariate normal distributions with means equal to the global mean fluorescence (top level of hierarchical model, Fig. 6A) and with a covariance matrix obtained by adding quadratically the variances from all layers of the hierarchical model. The custom R scripts used in the decoding analyses are publicly available on GitHub (https://github.com/giovannidiana/TRFS).

### Network model of *tph-1* and *daf-7* regulation

In this section we discuss our modelling approach for characterizing the regulatory interactions among neurons. As a short-hand notation we will denote the *ADF*, *ASI* and *NSM* neurons as *x*, *y* and *z* respectively. Here we introduce a computational method to characterize neuronal regulatory interactions based on the ratios between expression levels in mutants and wild type. We assigned three binary variables: *s_x_*, *s_y_* and *s_z_* to all neurons and then considered the 8 configurations generated by the triplet (*s_x_*, *s_y_*, *s_z_*) as in Fig S6C (right panel).

We embedded these 8 configurations into the cubic network depicted in Fig. S6C (left panel) allowing transitions between configurations that only differ by one of the binary variables. We then introduce parameters representing the level of interaction among neurons to define the transitions between configurations. The probabilities for each configuration can be interpreted as a representation of the circuit under a given choice of interaction parameters.

Given a transition rate matrix *W*, the probability vector of the 8 configurations *P* = {*p*_1_, ⋯, *p*_8_} satisfies the Master equation (*53*)

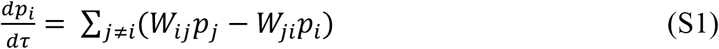

where the first term on the right hand side is a positive contribution due to incoming transitions from any node *j* connected to node *i* and a negative contribution due to outgoing transitions from node *i*. Note that the matrix element *W*_*ij*_ denotes the rate of the transition *j* → *i*. The steady state probabilities are then obtained by solving the linear equation 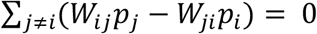.

To parameterize the transition rates, we introduced 9 parameters representing 3 self-regulations and 6 cross-regulations summarized by the regulatory matrix *w*

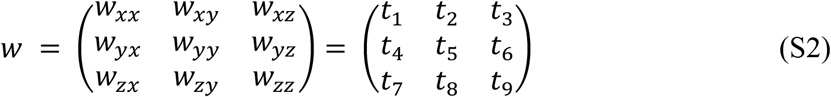

where each matrix element *w*_*αβ*_ characterizes the negative regulation of neuron *β* due to neuron *α*. To keep the analysis general, we assumed non-symmetric mutual regulation between neurons, i.e., *w*_*αβ*_ ≠ *w*_*βα*_. By using this regulatory matrix, we can parameterize the transition rate matrix *W* as

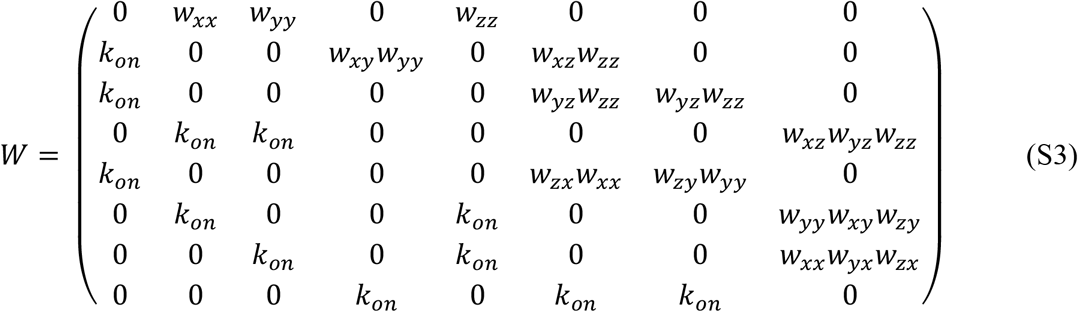

where *k*_*on*_is the rate at which each neuron switches to the ON state. For instance, let us consider the transition from configuration 2 to configuration 1 in Fig. S6C where ADF goes from an active to inactive state. The corresponding transition rate *W*_12_ = *w*_*xx*_reflects the role of *w*_*xx*_ in parameterizing the negative (self-)regulation of ADF. When *w*_*xx*_ is large, the transitions 2 *→* 1 are enhanced by the regulation, which reduces the (steady state) probability of ADF being active. Vice-versa, when *w*_*xx*_ is small, configuration 2 is more stable. Now, let us consider as a second example, the transition from configuration 4 = (1, 1, 0) where ADF and ASI are both ON, to configuration 2 = (1, 0, 0) where ASI has been switched OFF. The corresponding transition rate is *W*_24_ = *w*_*xy*_*w*_*yy*_. In this case the rate is affected by both ASI itself through the self-regulation term *w*_*yy*_ and ADF through *w*_*xy*_.

By setting the reference rate *k*_*on*_ = 1, all terms in the regulatory matrix that are larger than 1 generate down-regulation (in the case of cross-regulatory terms) or are smaller than *k*_*on*_ (in the case of self-regulatory terms).

Given a set of regulatory parameters *w*’s, we can obtain the probability of each configuration by considering the steady state of the Master equation dynamics. We can also derive the (marginal) probabilities of each neuron being in an active state from all configurations where that neuron is active:

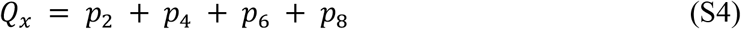

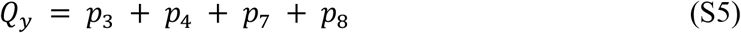

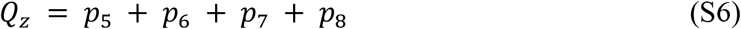

With this setting, we can mimic the effect of null mutant of a gene by removing from the network all edges arising from that gene. For instance, we can model the *daf-7*(*-*) mutant by removing edges *t*_4_, *t*_5_ and *t*_6_ as shown in Fig. S6D. Note that edges *t*_2_ and *t*_8_ are still in the network because in the *daf-7*(*-*) mutant the promoter is still present. The removal of an edge from the network is equivalent to setting the corresponding regulatory interaction *w* equal to 1. At fixed regulatory parameters, we can then compare how the probabilities of *ADF*, *ASI* and *NSM* being active change between wild-type and mutant networks. We can define the ratios between the probability of each neuron being active in wild type and each mutant condition as

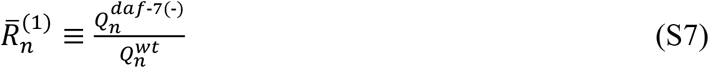

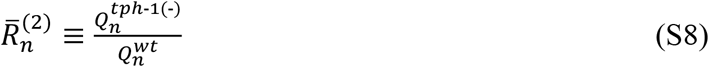

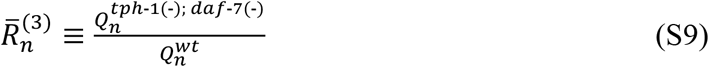

where *n* = *x, y, z*. To fit the regulatory interactions, we have constrained these quantities by using the same ratios obtained from the expression levels

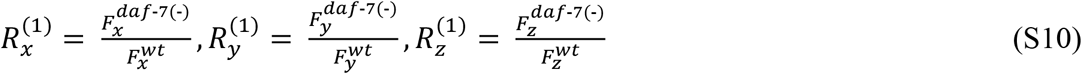

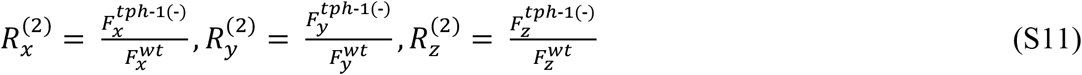

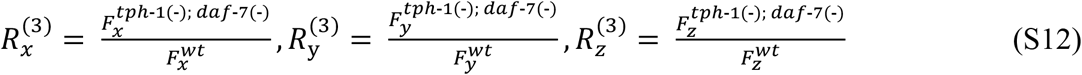

To estimate these ratios from the raw data, we paired imaging experiments performed on mutants with their corresponding wild type control, which were always performed on the same day. For all matching pairs, we estimated the mean expression levels for each neuron. We then obtained the average 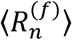 and the standard deviation 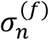 of the three mutant/wild-type ratios across all paired experiments.

To explore the parameter space, we employed a stochastic minimization of the cost function

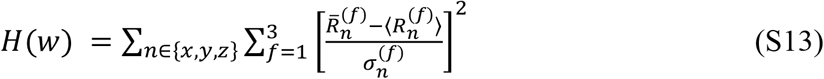

which depends on the set of regulatory parameters *w*. The algorithm proposes sequentially new parameters in the neighborhood of the previous ones (according to Euclidian distance) and accepts them if their corresponding cost function is reduced.

By performing this analysis, we found the sets of parameters that best represent the regulation among all three cells for each environmental condition of food and temperature tested (Fig. 4D). This result allowed us to identify regulations that are affected by temperature (Fig. 4D-E). We showed that these conclusions are robust in two ways. First, we used alternative methods of determining the network configurations. The same temperature-dependent regulations were identified when using the geometric means of the regulatory parameters weighted by goodness-of-fit, or when using the geometric means of the regulatory parameters for all models with a normalized likelihood greater than 95%. Second, the temperature-dependence did not arise by chance alone. In 100,0000 shuffles of the environmental conditions (i.e. shuffling the columns in Fig. 4D), monotonic temperature-dependent changes of the same magnitude for 4 edges occurred only in 14 permutations (Fig. 4E). Together, these computational analyses independently point to the key regulations that provide robustness to temperature.

## Acknowledgments

We thank the Horvitz Lab, the Bargmann Lab, and the *C. elegans* Genetics Center, which is funded by NIH Office of Research Infrastructure Programs (P40 OD010440) for reagents. We are grateful to C. Barnes and C. Houart for comments on this manuscript.

## Author Contributions

Designed experiments: all authors. Performed experiments: D.S.P., E.V.E., M.Z. Image processing: M.Z. Bayesian analysis and modelling: G.D. Data visualization: G.D., Q.C. Interpreted data: all authors. Wrote paper: Q.C., D.S.P., G.D., with input from all authors.

---

**Figure S1.**
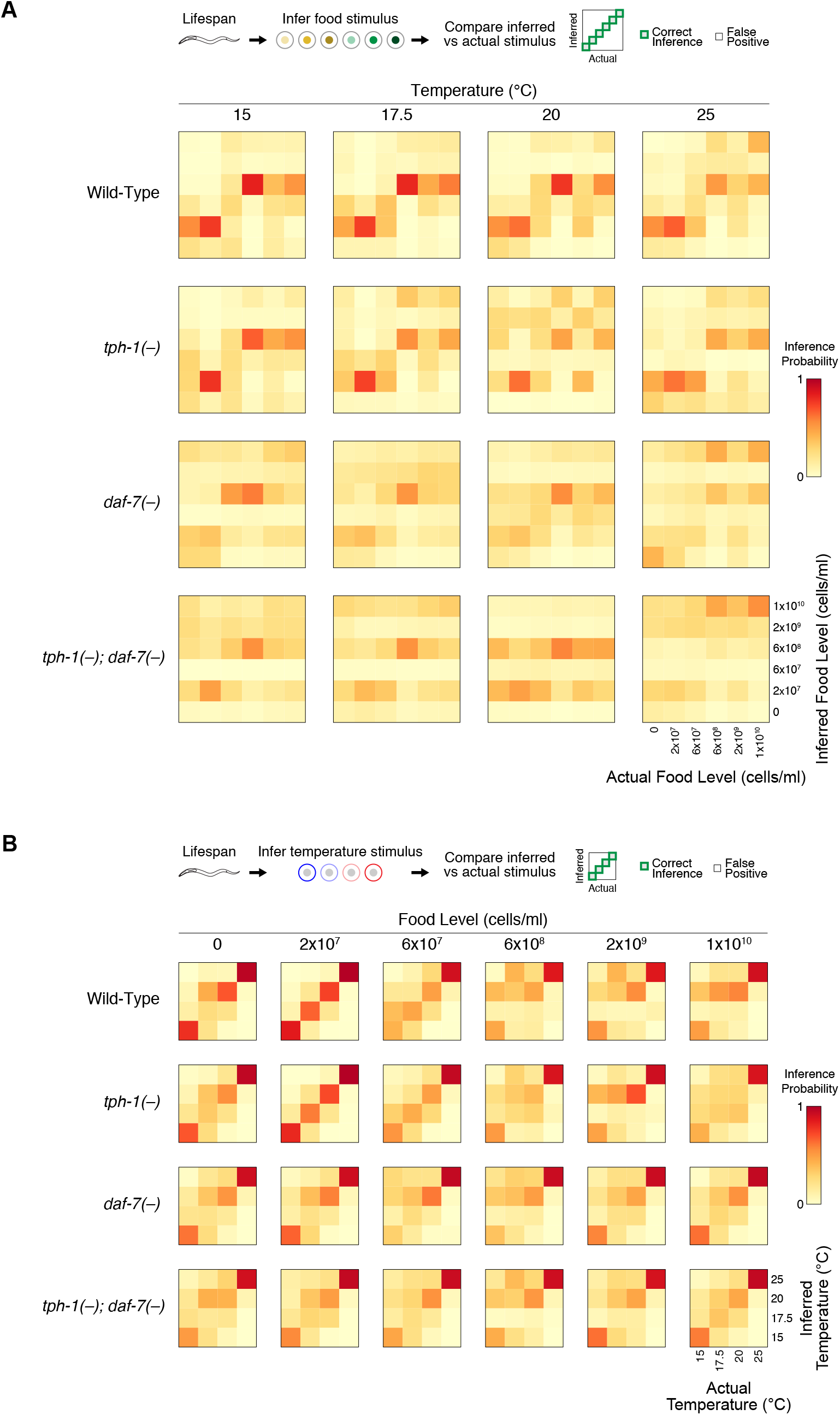
Accuracy of decoding food or temperature from lifespan is heterogenous and depends on *tph-1* and *daf-7*. **(A)** Top: summary of the food decoding process. Based on the lifespan of the individual, we inferred the food stimuli. The results summarized in a grid of heatmaps corresponding to food decoding for each temperature and genotype (bottom). Each heatmap indicates how frequently a food level was inferred, given the actual food stimulus. The diagonals represent the frequency of correct inferences, where the actual and inferred food levels are identical, whereas incorrect inferences are indicated by squares outside the diagonal. These heatmaps reveal that based on the lifespan responses, some food levels are well discriminated, whereas other food levels are not. For example, at 15 °C, wild-type animals can easily discriminate 2×10^7^ and 6×10^8^ bacterial cells/ml from other food levels, but tend to mistake no food for 2×10^7^ bacterial cells/ml. This inference pattern is stable in wild-type across temperature, only changing slightly at 25 °C, reflecting the robustness of food-sensing with temperature. By contrast, the inference patterns in the *tph-1*(*-*) and *daf-7*(*-*) mutants are not stable and changes more dramatically with temperature. **(B)** Top: summary of the temperature decoding process. Bottom: a grid of heatmaps depicting temperature inference at different food levels in different genotypes.

**Figure S2.**
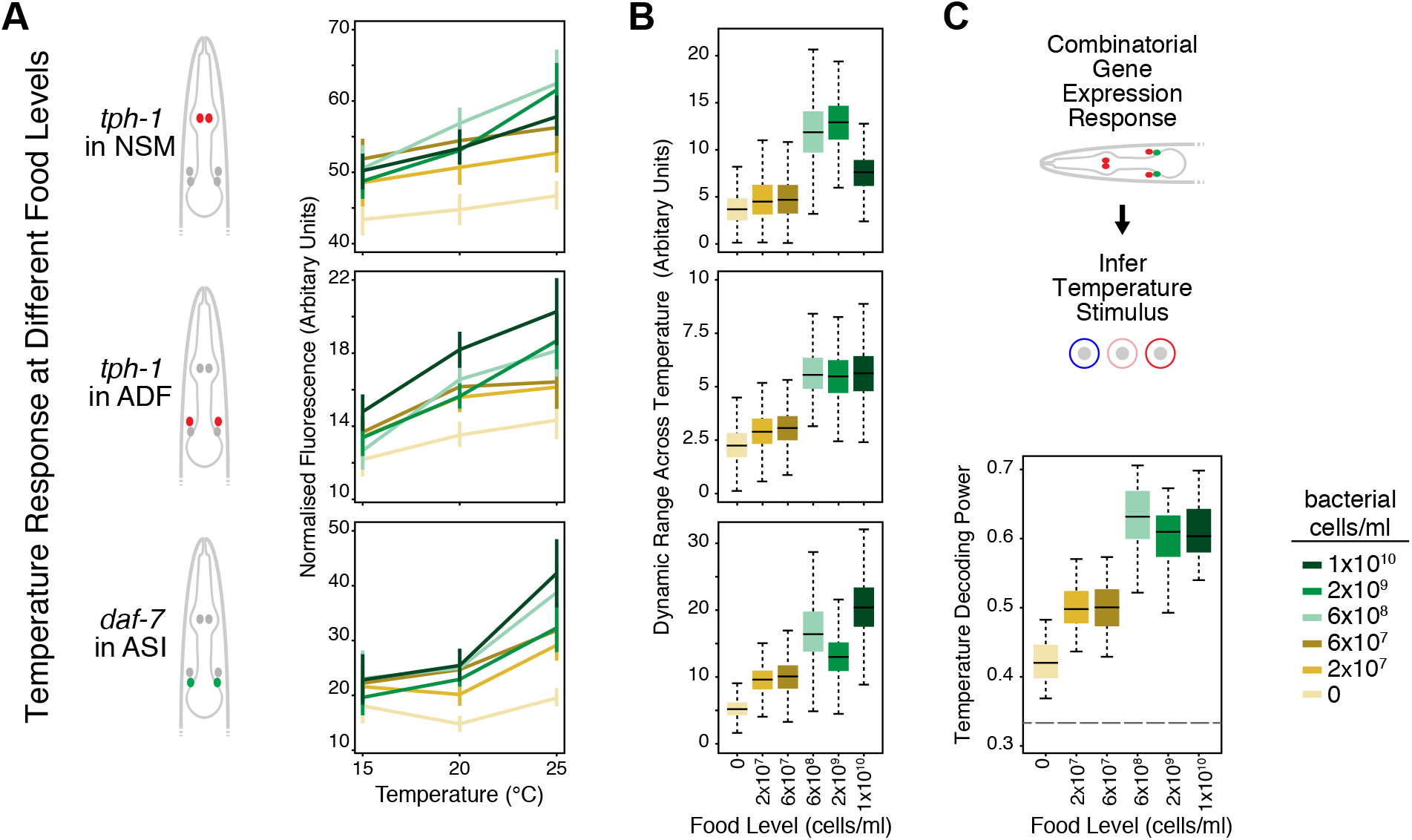
Effect of food on temperature responsiveness of *tph-1* and *daf-7* expression. **(A)** Wild-type expression of *tph-1* in NSM and ADF, and *daf-7* in ASI as a function of temperature. Each line corresponds to a different food level. See Figure 3b for legends. Diagrams on the left indicate the gene and cell. **(B)** Temperature-responsive dynamic range in the expression of *tph-1* in NSM and ADF, and *daf-7* in ASI at different food levels. **(C)** Wild-type temperature decoding power at different food levels based on the combinatorial expression of *tph-1* and *daf-7* in these three cells. Bayesian estimates are shown for all mean expression values and dynamic ranges (see Methods and Table S2). Dotted line in (C) indicates decoding power from random chance alone.

**Figure S3.**
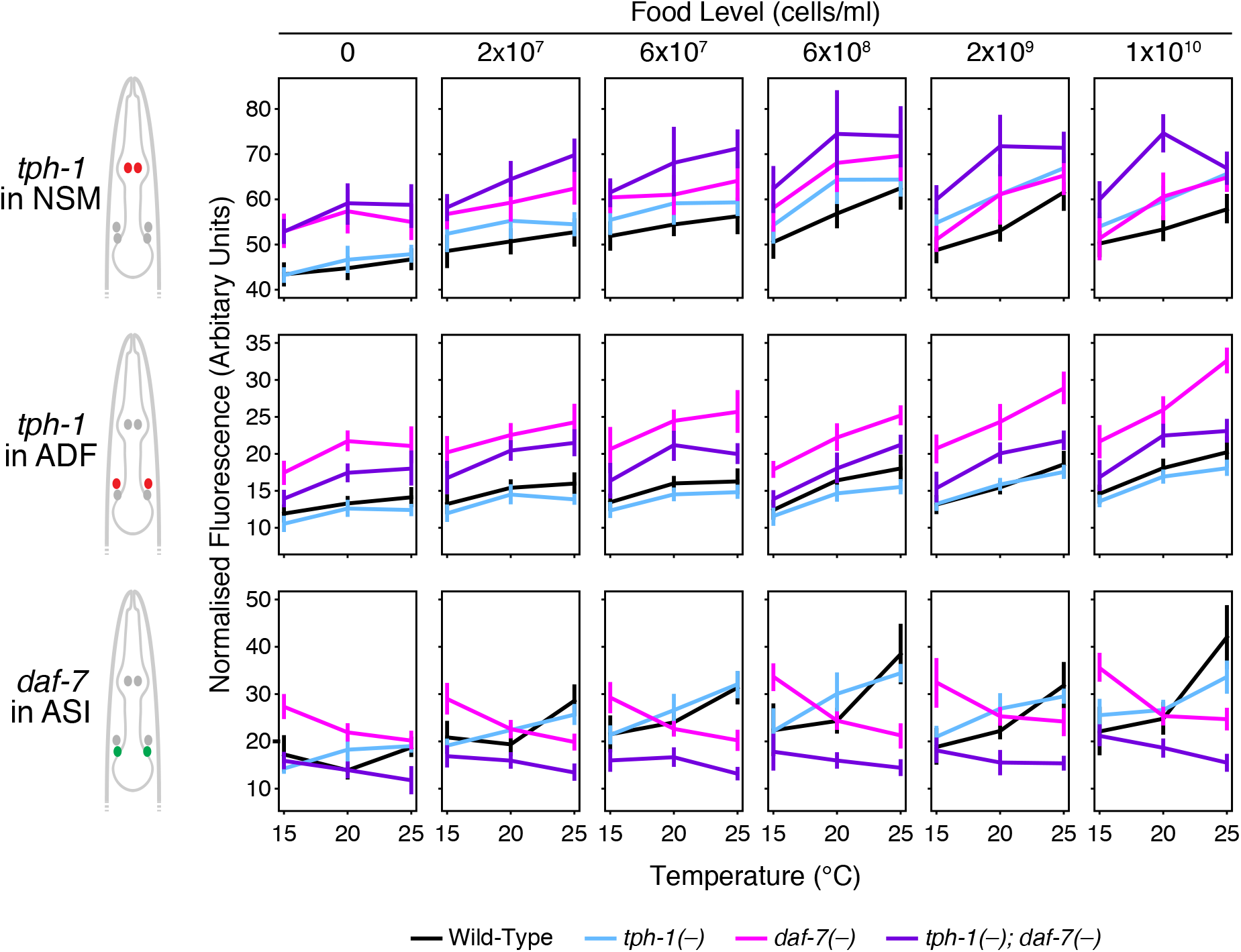
Effect of temperature on *tph-1* and *daf-7* expression as a function of food level. Bayesian estimates for expression values of *tph-1* in NSM and ADF, and *daf-7* in ASI are shown. Error bars denote 90% confidence intervals.

**Figure S4.**
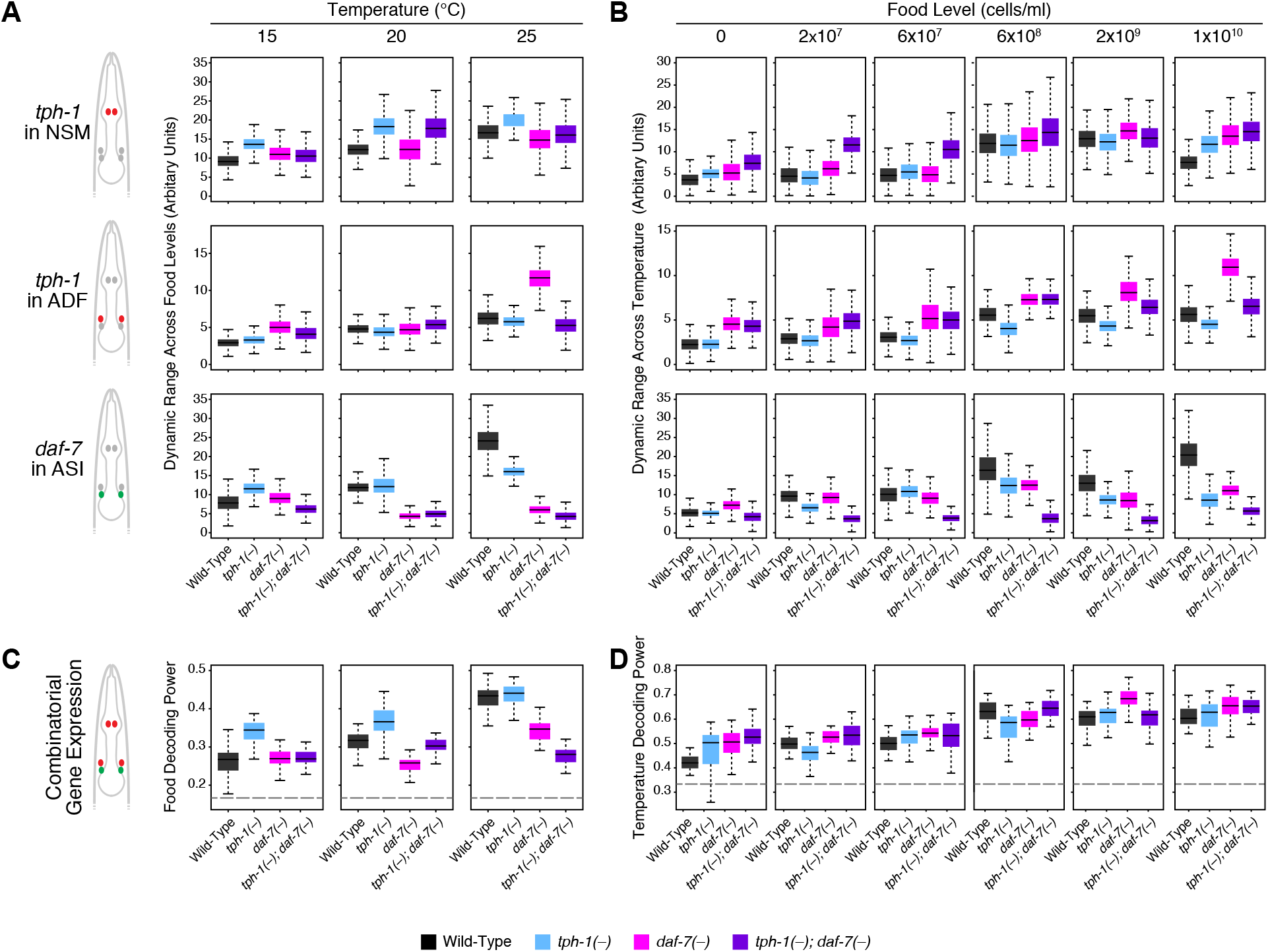
Effects of genotype, temperature, and food on dynamic range and decoding power of *tph-1* and *daf-7* expression. **(A)** Dynamic range of food-responsive *tph-1* expression in NSM and ADF, and *daf-7* expression in ASI at different temperatures. Diagrams on the left indicate the gene and cell. **(B)** Dynamic range of temperature-responsive *tph-1* expression in NSM and ADF, and *daf-7* expression in ASI at different food levels. **(C)** Food decoding power based on combinatorial expression of *tph-1* and *daf-7* in all three cells at each temperature. **(D)** Temperature decoding power based on combinatorial expression of *tph-1* and *daf-7* in all three cells at different food levels. (C-D) are derived from the decoding accuracies shown in Figure S5.

**Figure S5.**
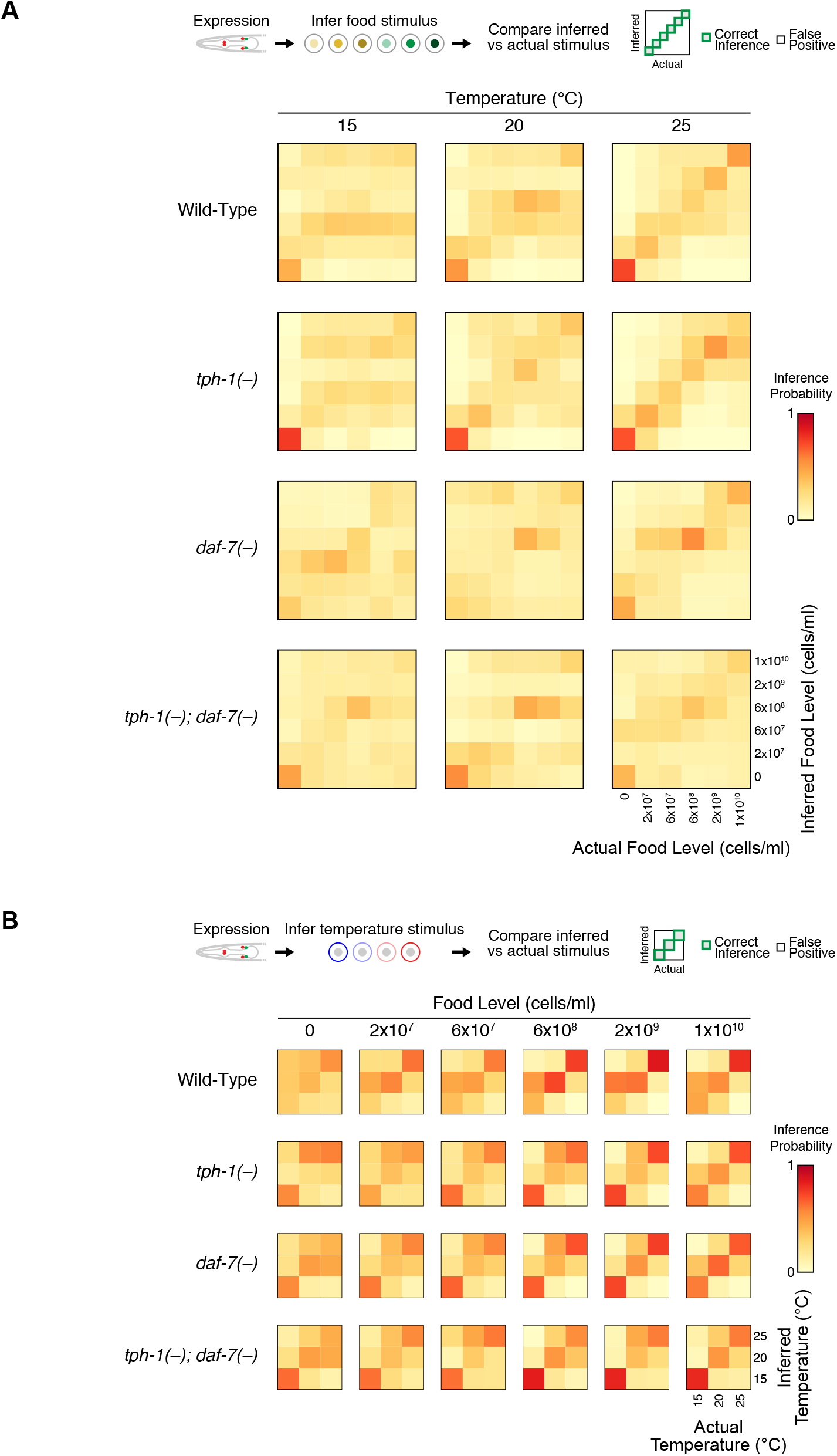
Accuracy of decoding food or temperature from gene expression. **(A)** Top: summary of the food decoding process using gene expression. Bottom: heatmaps depicting food decoding at different temperatures in different genotypes. **(B)** Top: summary of the temperature decoding process using gene expression. Bottom: heatmaps depicting temperatures decoding at different food levels in different genotypes.

**Figure S6.**
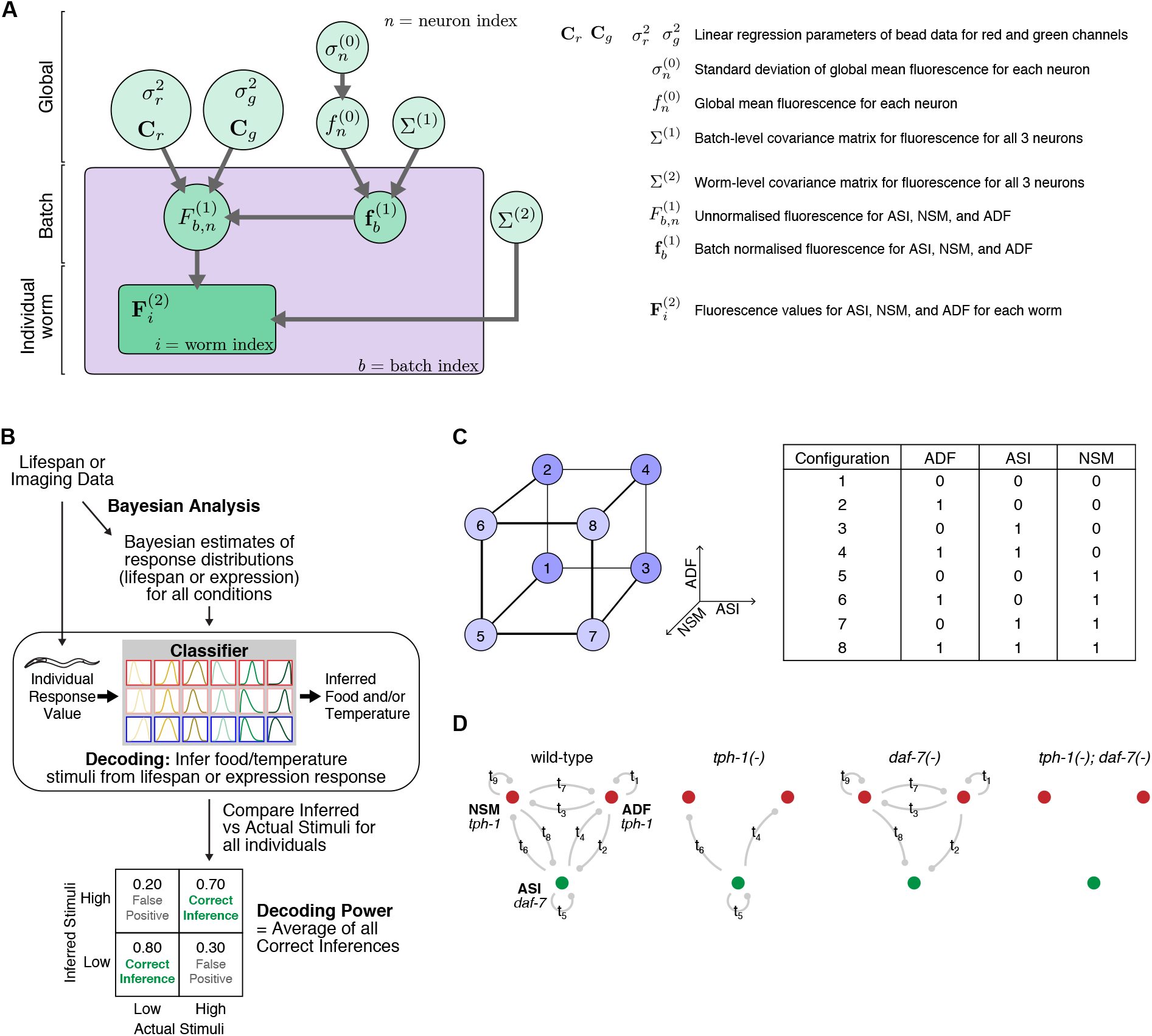
Bayesian analysis and network modelling. **(A)** Hierarchical Bayesian model for estimating the expression values of ASI, ADF, and NSM. **(B)** Schematic of the decoding analysis procedure. **(C)** Model representing possible states of ASI, ADF, and NSM and the transitions between these states (see supplementary methods for details). **(D)** The network is modelled with terms (*t_1_* to *t_9_*) for each directed cell-cell regulation. Mutants were represented in the network models as the absence of edges emanating from the cell normally expressing the mutated gene.

## References

1. M. A. Felix, M. Barkoulas, Pervasive robustness in biological systems. Nat Rev Genet 16, 483–496 (2015).

2. H. Kitano, Biological robustness. Nat Rev Genet 5, 826–837 (2004).

3. P. R. Hiesinger, B. A. Hassan, The Evolution of Variability and Robustness in Neural Development. Trends Neurosci 41, 577–586 (2018).

4. J. A. Mohawk, C. B. Green, J. S. Takahashi, Central and peripheral circadian clocks in mammals. Annu Rev Neurosci 35, 445–462 (2012).

5. E. Marder, M. L. Goeritz, A. G. Otopalik, Robust circuit rhythms in small circuits arise from variable circuit components and mechanisms. Curr Opin Neurobiol 31, 156–163 (2015).

6. B. Conti, Considerations on temperature, longevity and aging. Cell Mol Life Sci 65, 1626–1630 (2008).

7. J. Alcedo, W. Maier, Q. Ch’ng, Sensory influence on homeostasis and lifespan: molecules and circuits. Advances in experimental medicine and biology 694, 197–210 (2010).

8. L. Fontana, L. Partridge, Promoting health and longevity through diet: from model organisms to humans. Cell 161, 106–118 (2015).

9. W. Mair, P. Goymer, S. D. Pletcher, L. Partridge, Demography of dietary restriction and death in Drosophila. Science 301, 1731–1733 (2003).

10. N. M. Templeman, C. T. Murphy, Regulation of reproduction and longevity by nutrient-sensing pathways. J Cell Biol 217, 93–106 (2018).

11. N. A. Bishop, L. Guarente, Genetic links between diet and lifespan: shared mechanisms from yeast to humans. Nature reviews Genetics 8, 835–844 (2007).

12. S. J. Lee, C. Kenyon, Regulation of the longevity response to temperature by thermosensory neurons in Caenorhabditis elegans. Curr Biol 19, 715–722 (2009).

13. R. Xiao et al., A genetic program promotes C. elegans longevity at cold temperatures via a thermosensitive TRP channel. Cell 152, 806–817 (2013).

14. E. L. Greer, A. Brunet, Different dietary restriction regimens extend lifespan by both independent and overlapping genetic pathways in C. elegans. Aging cell 8, 113–127 (2009).

15. P. Kapahi, M. Kaeberlein, M. Hansen, Dietary restriction and lifespan: Lessons from invertebrate models. Ageing Res Rev 39, 3–14 (2017).

16. C. E. Riera, A. Dillin, Emerging Role of Sensory Perception in Aging and Metabolism. Trends Endocrinol Metab 27, 294–303 (2016).

17. N. Stroustrup et al., The temporal scaling of Caenorhabditis elegans ageing. Nature 530, 103–107 (2016).

18. E. V. Entchev et al., A gene-expression-based neural code for food abundance that modulates lifespan. Elife 4, e06259 (2015).

19. D. S. Patel et al., Quantification of Information Encoded by Gene Expression Levels During Lifespan Modulation Under Broad-range Dietary Restriction in C. elegans. J Vis Exp, (2017).

20. H. Schulenburg, M. A. Felix, The Natural Biotic Environment of Caenorhabditis elegans. Genetics 206, 55–86 (2017).

21. P. Dayan, L. F. Abbott, Theoretical Neuroscience: Computational And Mathematical Modeling of Neural Systems. (Massachusetts Institute of Technology Press, 2005).

22. A. A. Granados et al., Distributed and dynamic intracellular organization of extracellular information. Proc Natl Acad Sci U S A 115, 6088–6093 (2018).

23. P. Ren et al., Control of C. elegans larval development by neuronal expression of a TGF-beta homolog. Science 274, 1389–1391 (1996).

24. Y. Zhang, H. Lu, C. I. Bargmann, Pathogenic bacteria induce aversive olfactory learning in Caenorhabditis elegans. Nature 438, 179–184 (2005).

25. J. Y. Sze, M. Victor, C. Loer, Y. Shi, G. Ruvkun, Food and metabolic signalling defects in a Caenorhabditis elegans serotonin-synthesis mutant. Nature 403, 560–564 (2000).

26. W. M. Shaw, S. Luo, J. Landis, J. Ashraf, C. T. Murphy, The C. elegans TGF-beta Dauer pathway regulates longevity via insulin signaling. Curr Biol 17, 1635–1645 (2007).

27. K. Ashrafi, Obesity and the regulation of fat metabolism. WormBook: the online review of C elegans biology, 1–20 (2007).

28. A. J. Chang, N. Chronis, D. S. Karow, M. A. Marletta, C. I. Bargmann, A distributed chemosensory circuit for oxygen preference in C. elegans. PLoS Biol 4, e274 (2006).

29. M. L. Brown, A. L. Schneyer, Emerging roles for the TGFbeta family in pancreatic beta-cell homeostasis. Trends in endocrinology and metabolism: TEM 21, 441–448 (2010).

30. F. Oury, G. Karsenty, Towards a serotonin-dependent leptin roadmap in the brain. Trends in endocrinology and metabolism: TEM 22, 382–387 (2011).

31. G. Diana et al., Genetic control of encoding strategy in a food-sensing neural circuit. Elife 6, (2017).

32. W. S. Schackwitz, T. Inoue, J. H. Thomas, Chemosensory neurons function in parallel to mediate a pheromone response in C. elegans. Neuron 17, 719–728 (1996).

33. M. Fletcher, D. H. Kim, Age-Dependent Neuroendocrine Signaling from Sensory Neurons Modulates the Effect of Dietary Restriction on Longevity of Caenorhabditis elegans. PLoS Genet 13, e1006544 (2017).

34. T. Gallagher, J. Kim, M. Oldenbroek, R. Kerr, Y. J. You, ASI regulates satiety quiescence in C. elegans. J Neurosci 33, 9716–9724 (2013).

35. J. L. Rhoades et al., ASICs Mediate Food Responses in an Enteric Serotonergic Neuron that Controls Foraging Behaviors. Cell 176, 85–97 e14 (2019).

36. A. Zaslaver et al., Hierarchical sparse coding in the sensory system of Caenorhabditis elegans. Proc Natl Acad Sci U S A 112, 1185–1189 (2015).

37. K. Chung, M. M. Crane, H. Lu, Automated on-chip rapid microscopy, phenotyping and sorting of C. elegans. Nat Methods 5, 637–643 (2008).

38. M. Zhan et al., Automated Processing of Imaging Data through Multi-tiered Classification of Biological Structures Illustrated Using Caenorhabditis elegans. PLoS Comput Biol 11, e1004194 (2015).

39. G. Angelo, M. R. Van Gilst, Starvation protects germline stem cells and extends reproductive longevity in C. elegans. Science 326, 954–958 (2009).

40. T. O’Leary, E. Marder, Temperature-Robust Neural Function from Activity-Dependent Ion Channel Regulation. Curr Biol 26, 2935–2941 (2016).

41. A. Millius, H. R. Ueda, Systems Biology-Derived Discoveries of Intrinsic Clocks. Front Neurol 8, 25 (2017).

42. G. M. Edelman, J. A. Gally, Degeneracy and complexity in biological systems. Proc Natl Acad Sci U S A 98, 13763–13768 (2001).

43. M. Beverly, S. Anbil, P. Sengupta, Degeneracy and neuromodulation among thermosensory neurons contribute to robust thermosensory behaviors in Caenorhabditis elegans. J Neurosci 31, 11718–11727 (2011).

44. E. C. Cropper, A. M. Dacks, K. R. Weiss, Consequences of degeneracy in network function. Curr Opin Neurobiol 41, 62–67 (2016).

45. S. A. Haddad, E. Marder, Circuit Robustness to Temperature Perturbation Is Altered by Neuromodulators. Neuron 100, 609–623 e603 (2018).

46. T. Stiernagle, Maintenance of C. elegans. WormBook, 1–11 (2006).

47. K. C. Cheng, R. Klancer, A. Singson, G. Seydoux, Regulation of MBK-2/DYRK by CDK-1 and the pseudophosphatases EGG-4 and EGG-5 during the oocyte-to-embryo transition. Cell 139, 560–572 (2009).

48. J. M. Parry et al., EGG-4 and EGG-5 Link Events of the Oocyte-to-Embryo Transition with Meiotic Progression in C. elegans. Curr Biol 19, 1752–1757 (2009).

49. M. M. Crane et al., Autonomous screening of C. elegans identifies genes implicated in synaptogenesis. Nat Methods 9, 977–980 (2012).

50. M. A. Unger, H. P. Chou, T. Thorsen, A. Scherer, S. R. Quake, Monolithic microfabricated valves and pumps by multilayer soft lithography. Science 288, 113–116 (2000).

51. M. Plummer, rjags: Bayesian Graphical Models using MCMC. https://CRAN.R-project.org/package=rjags, (2018).

52. B. P. Carlin, T. A. Louis, B. P. Carlin, Bayesian methods for data analysis. Chapman & Hall/CRC texts in statistical science series (CRC Press, Boca Raton, ed. 3rd, 2009).

53. N. G. van Kampen, Stochastic processes in physics and chemistry. North-Holland personal library (Elsevier, Amsterdam; Boston, ed. 3rd, 2007).

